# Tear Proteomics Reveals RAGE and NLRP3 Inflammasome Pathway Activation in Lacrimal Glands of a Sjögren’s Disease Mouse Model

**DOI:** 10.64898/2026.07.15.738677

**Authors:** Xiaoyang Li, Mario Alba, Daniel J. Fernandez, Alison V. Ramirez, Sara Abdelhamid, Chenyi Wang, Maria C. Edman, Julian Whitelegge, Sarah F. Hamm-Alvarez

**Affiliations:** Department of Ophthalmology, University of Southern California Keck School of Medicine, Los Angeles, California, United States; Department of Pharmacology and Pharmaceutical Sciences, Alfred E. Mann School of Pharmacy, University of Southern California, Los Angeles, California, United States; Pasarow Mass Spectrometry Laboratory and Neuropsychiatric Institute-Semel Institute for Neuroscience and Human Behavior, David Geffen School of Medicine, University of California at Los Angeles, Los Angeles, California, United States

## Abstract

**Purpose:** To characterize tear proteome changes in male non-obese diabetic (NOD) mice with Sjögren’s disease (SjD)-like autoimmune dacryoadenitis and determine whether identified tear proteins are associated with lacrimal gland (LG) pathogenesis.

**Methods:** Tears were collected from 14-week-old male NOD mice and age-matched male BALB/c controls and analyzed by tandem mass tag (TMT)-based liquid chromatography tandem mass spectrometry (LC-MS/MS). Differentially expressed proteins (DEPs) were defined using adjusted *P* < 0.05 and absolute fold change > 1.5. Functional enrichment analysis was performed using Enrichr. Selected upregulated DEPs were further examined in tear and LG samples from independent mouse cohorts using Western blotting, immunofluorescence, and RT-qPCR.

**Results:** A total of 142 proteins were quantified across all tear samples. Hierarchical clustering and principal component analysis (PCA) separated NOD from BALB/c tear proteomes. A total of 41 proteins were differentially expressed in NOD mouse tears (33 increased, 8 decreased). Increased tear proteins were enriched in immune and inflammatory responses, secretory compartments, RAGE receptor binding, glutathione metabolism, oxidative stress, and redox regulation. S100A8/A9, GSTO1-1, Gal-3, and pIgR/secretory component (SC) were increased in both NOD mouse tears and LG. In NOD LG, increased S100A8/A9 was accompanied by elevated RAGE, whereas increased GSTO1-1 was associated with increased NLRP3, cleaved caspase-1, cleaved gasdermin, cleaved IL-1β, and increased *Il1b*, *Il18*, and *Il18r* gene expression.

**Conclusions:** Male NOD mouse tears contain disease-related proteins reflecting pathological inflammatory and epithelial changes in the LG including increased RAGE signaling, NLRP3 inflammasome activation, and altered epithelial transcytosis.

## INTRODUCTION

Sjögren’s disease (SjD) is a chronic autoimmune disorder characterized by progressive exocrinopathies of the lacrimal gland (LG) and salivary gland (SG) associated with lymphocytic infiltration^1^. These infiltrates initiate and perpetuate a cascade of immune and inflammatory events that disrupt epithelial cell integrity and impair glandular secretory functions, leading to the hallmark sicca symptoms of dry eye and dry mouth as well as development of systemic symptoms^1^. Despite the significant clinical burden associated with ocular SjD^2^, current diagnostic approaches rely on assessment of subjective symptoms which are shared across other types of dry eye disease (DED), in combination with invasive or relatively insensitive clinical tests^3–5^, leaving many patients undiagnosed. Existing therapies largely provide topical symptomatic relief and fail to suppress LG inflammation and/or restore its normal secretory functions^6,7^. Although advances have been made in defining dysregulated immune profiles in glandular infiltrates and in circulation^8,9^, the molecular mechanisms that sustain SjD-associated autoimmune dacryoadenitis remain incompletely understood^10^.

The tear film supplies proteins to the ocular surface system that are critical for supporting its health and protecting it from pathogens^11,12^. Under physiological conditions, LG acinar cells (LGAC) produce many of these proteins through 1) tear protein production and secretion via regulated exocytosis and 2) transcytosis of proteins from the basolateral interstitium, a process including polymeric immunoglobulin receptor (pIgR)-mediated transport of dimeric IgA^13–16^. Chronic inflammation disrupts these secretory functions, altering tear composition and local immune responses^17–19^. In parallel, epithelial stress and injury within the inflamed LG can elicit release of damage-associated molecular pattern molecules (DAMPs), increase oxidative stress, and contribute to chronic inflammatory remodeling of glandular tissues, all of which may engage innate immune signaling pathways^8,20–23^. When activated, these processes may amplify inflammatory responses and contribute to progressive autoimmune exocrinopathy^24,25^. While other tissues and organs of the ocular surface system also contribute proteins and other components to tears, the major contributions of the LG to tear protein production suggest that investigation of changes in the tear proteome in SjD will provide insights into LG disease pathology.

Among the innate immune response pathways that can propagate inflammation, the NOD-like receptor pyrin domain containing 3 (NLRP3) inflammasome signaling has attracted interest^26–28^. The NLRP3 inflammasome coordinates caspase-1 activation, maturation of interleukin-1β and -18 (IL-1β and IL-18), and cleavage of gasdermin D (GSDMD), ultimately leading to release of pro-inflammatory cytokines and induction of a regulated cell death pathway known as pyroptosis^28^. Oxidative stress, a recognized feature of autoimmune exocrinopathy in SjD^29^, can regulate the activation of the NLRP3 inflammasome under conditions of sustained inflammation^28^. Although NLRP3 inflammasome signaling has been implicated in DED and exocrine gland dysfunction, its activation and specific contribution to LG pathology in SjD remains incompletely characterized^26–28,30^.

The Non-Obese Diabetic (NOD) mouse is one of the most thoroughly characterized models for investigating SjD-related exocrinopathies, with the males spontaneously developing ocular symptoms of SjD^18,31–37^. Male NOD mice exhibit multiple hallmarks of the autoimmune dacryoadenitis that characterizes human SjD including reduced tear secretion^31^, lymphocytic infiltration of LG including the formation of tertiary lymphoid structures^37^, changes in LG extracellular matrix^32^, and the development of tear-specific biomarkers of SjD^19,31,36^. As previously published^18,36,37^, we use the male BALB/c mouse as a healthy sex-and age-matched control strain.

Tear fluid is a readily accessible biofluid that contains molecular signatures linked to LG health^12,38^ as well as molecular signatures related to inflammation and stress responses in disease^39^. Quantitative LC–MS/MS-based proteomics serves as a platform for multiplexed, comprehensive profiling of disease-associated proteome alterations, enabling the discovery of candidate biomarkers, and potential new therapeutic targets. The present study leverages quantitative proteomics using tears from the male NOD mouse model of SjD-like autoimmune dacryoadenitis to 1) characterize tear proteome alterations potentially associated with SjD and 2) to link changes in differentially expressed tear proteins to development of LG pathology.

## MATERIALS AND METHODS

### Animals

14-week-old male NOD (#001976) mice were used as a model for SjD-like autoimmune dacryoadenitis while age-matched male BALB/c (#000651) mice served as healthy controls. Both strains were purchased from The Jackson Laboratory (Bar Harbor, ME, USA). All procedures conducted on animals were approved by the University of Southern California’s Institutional Animal Care and Use Committee (IACUC) and conducted in accordance with the Guide for the Care and Use of Laboratory Animals and the ARVO Statement for the Use of Animals in Ophthalmic and Vision Research.

### Tear and Tissue Collection

Mice were anesthetized with Ketamine (60-70 mg/kg)/Xylazine (5-10 mg/kg). Stimulated tears were collected using glass capillaries as previously described^35^, and flash-frozen until use. Samples were acquired from two separate cohorts of mice: a discovery cohort providing tears for quantitative proteomics and a validation cohort providing additional tears. For validation of changes in LG protein expression by Western blotting, separate cohorts of mice were euthanized, and unstimulated LG were collected and processed for protein extraction, immunofluorescence or RT-qPCR.

### Quantitative Proteomics Sample Processing

Total protein in tear samples from NOD and BALB/c mice (n = 4 samples/strain with each sample comprised of pooled tears from 5 mice) was quantified using the Micro BCA Protein Assay Kit (Thermo Fisher Scientific, Waltham, MA, USA). 300 µg of tear protein from each sample was processed in lysis buffer containing 12 mM sodium lauroyl sarcosine, 0.5% sodium deoxycholate, and 50 mM triethylammonium bicarbonate (TEAB), supplemented with Halt^TM^ Protease and Phosphatase Inhibitor Cocktail (Thermo Fisher Scientific, Waltham, MA). Samples were sonicated for 10 min, reduced with 10 mM tris(2-carboxyethyl) phosphine (TCEP) and alkylated with 40 mM chloroacetamide for 10 min at 95°C, and then diluted in 50 mM TEAB and digested overnight with Sequencing Grade Modified Trypsin (Promega, Madison, WI, USA). Peptides were extracted using ethyl acetate/trifluoracetic acid (TFA, 100:1, v/v), with thorough vortexing (5 min) and centrifugation (16,000 × g, 5 min), then dried in a centrifugal vacuum concentrator. Peptides were then desalted using Pierce™ Peptide Desalting Spin Columns (Thermo Fisher Scientific, Waltham, MA) per the manufacturer’s protocol. The collected peptides were dried and quantified using Pierce™ Quantitative Peptide Assays (Thermo Fisher Scientific, Waltham, MA). Peptides were then chemically labeled with TMTpro™ 16plex Label Reagent Set (Thermo Fisher Scientific, Waltham, MA) according to the manufacturer’s protocol. The samples were pooled together and desalted again using Pierce™ Peptide Desalting Spin Columns (Thermo Fisher Scientific, Waltham, MA) per the manufacturer’s protocol. The collected eluents were dried, reconstituted in acetonitrile/water/formic acid (2:98:0.1, v/v/v), and fractionated using the Pierce™ High pH Reversed-Phase Peptide Fractionation Kit (Thermo Fisher Scientific, Waltham, MA) according to the manufacturer’s protocol.

### LC-MS/MS Acquisition

Peptide fractions (5 µL) were injected onto a reverse-phase nanobore HPLC column (QuanDx, San Jose, CA, USA; C18, 1.8 μm particle size, 360 μm × 20 cm, 150 μm ID) equilibrated in solvent A (0.1% formic acid) and separated at 300 nL/min on an Easy-nLC 1200 system (Thermo Fisher Scientific, Waltham, MA) using a solvent B gradient (acetonitrile/water/formic acid, 98/2/0.1, v/v/v) (min/%B: 0/0, 5/3, 18/7, 74/12, 144/24, 153/27, 162/40, 164/80, 174/80, 176/0, 180/0). The column effluent was directed to a nanospray ionization source coupled to a Q Exactive Plus hybrid quadrupole-Orbitrap mass spectrometer (Thermo Fisher Scientific, Waltham, MA) operated in positive-ion, data-dependent acquisition mode. Full MS scans were acquired over m/z 350–1700 (AGC target 3 × 10^6^; maximum injection time 100 ms; Full Width at Half Maximum (FWHM) resolution 70,000 at m/z 200) and up to 15 MS/MS scans (quadrupole isolation of charge states 2-7, isolation window 1.5 m/z) with previously optimized fragmentation conditions (normalized collision energy of 28, AGC target 1 x 10^5^, 100 ms maximum injection time, FWHM resolution 17,500 at m/z 200).

### Proteomics Data Processing and Protein Quantification

Raw files from all fractions were processed in Proteome Discoverer™ (v2.3, Thermo Fisher Scientific, Waltham, MA). Database searching was performed against the UniProt mouse reviewed protein database using the SEQUEST-HT search engine. A reversed-sequence decoy strategy was used to control false discovery rate, with an FDR threshold of <1% for high-confidence peptide identification. Tryptic peptides unique to individual proteins were used for protein identification and relative quantification across samples. Only proteins with valid quantitative measurements across all analyzed samples were retained for downstream statistical and bioinformatic analyses.

### Functional Enrichment Analyses

Functional enrichment analyses were performed using Enrichr^40,41^. Differentially expressed proteins (DEPs) were defined using an adjusted *P* < 0.05 and an absolute fold change (FC) > 1.5 and analyzed separately for proteins increased and decreased in NOD tears compared to healthy control BALB/c mice. Gene Ontology (GO) enrichment analysis was conducted across the Biological Process (BP), Molecular Function (MF), and Cellular Component (CC) categories^42^. In parallel, pathway enrichment analysis was performed using the WikiPathways Mouse database^43^. Enrichment significance was calculated in Enrichr using a Fisher’s exact test comparing the input DEP list to the corresponding background database gene set. For each enriched term, Enrichr provides a nominal *P*, a Benjamini–Hochberg adjusted *P*, a z-score reflecting deviation from expected rank, and a Combined Score defined as ln(*P*) × z-score, where *P* represents the nominal *P* value^40,41,44^. For visualization purposes, enrichment results were ranked based on Combined Score and displayed as log10(Combined Score). Representative enriched terms were selected for presentation based on statistical significance and relevance to disease pathology; corresponding nominal p-values were displayed alongside each term.

### Western Blotting

Collected tears (n = 4 samples/strain; each sample comprised tears pooled from 3 mice) and LG samples (n = 4 samples/strain; one LG per sample) were lysed in RIPA buffer containing protease/phosphatase inhibitor cocktail. (Cell Signaling Technology, Danvers, MA). Protein concentration was measured using the Micro BCA Protein Assay Kit (Thermo Fisher Scientific, Waltham, MA). Samples were spiked with 4X Laemmli SDS sample buffer (Bio-Rad, Hercules, CA, USA) containing β-mercaptoethanol for 5 min at 95°C and then 40 μg of total protein was loaded on Tris-Glycine gels (Thermo Fisher Scientific, Waltham, MA) before electrophoresis. Proteins on gels were transferred to nitrocellulose membranes (Thermo Fisher Scientific, Waltham, MA) and stained for total protein (Li-Cor, Lincoln, NE, USA) to enable normalization to protein loading by signal intensity. Membranes were incubated in blocking buffer (Rockland, Limerick, PA, USA), and then incubated overnight in primary antibodies at the desired dilution at 4°C. After this, membranes were washed with Tris-buffered saline supplemented with 0.2% Tween (TBST) and incubated with 1:2000 diluted IR680 secondary antibodies (Li-Cor, Lincoln, NE) at room temperature for 1 h. After TBST washes, membranes were imaged with an Odyssey Licor imaging system. All primary and secondary antibodies used for Western blotting are listed in **Supplemental Table 1**, along with their relevant dilutions.

### LG Immunofluorescence

Unstimulated LGs were collected and fixed in 4% paraformaldehyde and 4% sucrose for 3 h at room temperature (RT) followed by 30% sucrose at 4°C overnight before embedding in Optimal Cutting Temperature (OCT) compound (VWR, Radnor, PA, USA). OCT-embedded LG were cut into 5 μm thin sections and mounted on Superfrost Plus microslides (VWR, Radnor, PA). The cryosections were quenched with 50 mM NH_4C_l in PBS for 20 min and permeabilized with 0.3% Triton X-100 for 30 min. Sections were blocked in 5% BSA with 0.3% Triton X-100 for 3 h at RT and incubated with the primary antibodies listed in **Supplemental Table 1** at either 1:100 or 1:200 dilution overnight at 4°C. After three washes in PBS, slides were incubated with secondary antibodies listed in **Supplemental Table 1**, as well as DAPI (Invitrogen, Carlsbad, CA, USA) and Rhodamine-phalloidin (Invitrogen, Carlsbad, CA) for 1 h at 37°C, washed three times in PBS and cover-slipped after application of 1 drop of Prolong Gold Antifade mounting medium (Invitrogen, Carlsbad, CA), and finally imaged with a Zeiss 800 LSM confocal fluorescence microscopy equipped with Airyscan processing (Zeiss, Thornwood, NY, USA), using either a 63× oil 1.4 NA objective or a 20× 0.8 NA objective. Images were processed equivalently across matched groups, with the brightness and contrast settings for each protein of interest kept consistent, using ZEN (Zeiss, Thornwood, NY), QuPath v0.5.1^45^, and Sketch (Sketch B.V., Eindhoven, the Netherlands) softwares.

### RT-qPCR

Total RNA from unstimulated LG of male NOD and BALB/c mice was extracted using the RNeasy plus Universal Mini Kit (Qiagen, Hilden, Germany) according to the manufacturer’s instructions. The concentration and purity of RNA was assessed using a Nanodrop. cDNA templates were synthesized from 4 µg RNA using TaqMan® gene expression. RT-qPCR was performed using TaqMan minor groove binder probes for *Il1b* (Mm00434228_m1), *Il18* (Mm00434225_m1), and *Il18r* (Mm00515180_m1). All primers were from Thermo Fisher Scientific (Waltham, MA). Gene expression levels were normalized to a housekeeping gene, GAPDH, and calculated using the ΔΔCt method.

### Statistical Analysis

All statistical analyses were performed using GraphPad Prism (Boston, MA, USA). For proteomic differential expression analysis, adjusted *P* values were calculated using the Benjamini–Hochberg method to control the false discovery rate. DEPs were defined as adjusted *P* < 0.05 and an absolute FC > 1.5. For Western blot and qPCR comparisons between BALB/c and NOD mice, statistical significance was assessed using the two-tailed Mann–Whitney U test. Data are presented as median ± interquartile range (IQR) unless otherwise specified. Each dot represents one biological replicate. A *P* < 0.05 was considered statistically significant.

## RESULTS

### Global proteome differences are seen between tears from male NOD and BALB/c mice

Scheme 1 shows the workflow for the tear proteomics analysis. A total of 142 proteins were quantified in all analyzed samples using Tandem Mass Tag (TMT)-based quantitative proteomics with a false discovery rate (FDR) of 1%. To investigate the differences in the tear proteomes of BALB/c and NOD mice, a hierarchical clustering analysis was performed, with results visualized in a heatmap (**Fig. 1A**). The heatmap showed clear clustering of the tear samples from healthy control BALB/c mice versus NOD mice and illustrated standardized relative protein abundance patterns across individual samples. This distinct clustering pattern was further supported by principal component analysis (PCA; **Fig. 1B**) which revealed distinct separation between the tear proteomes of BALB/c and NOD mice, with PC1 and PC2 explaining 61.9% and 9.9% of the total variance, respectively. A volcano plot was generated to visualize DEPs with an absolute FC > 1.5, and an adjusted *P* < 0.05 (**Fig. 1C**). A total of 41 DEPs were found in NOD mouse tears, including 33 upregulated and 8 downregulated proteins.

**Scheme 1.**
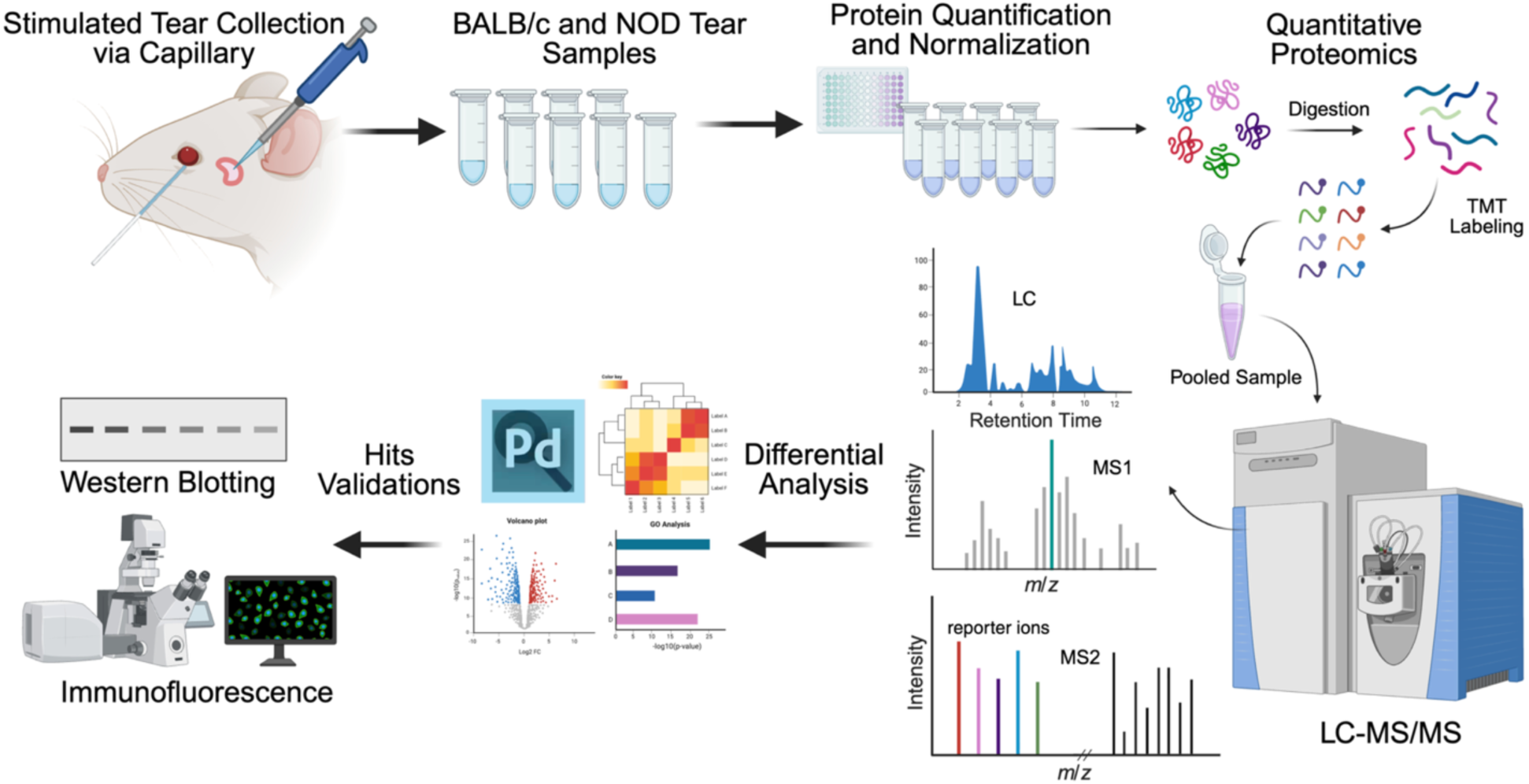
Tear proteomics study workflow. Tears were collected from 14-week-old male non-obese diabetic (NOD) mice as well as age-matched healthy control BALB/c male mice (n = 4 per strain, each sample contained pooled tears from 5 mice) using capillaries following topical carbachol stimulation of the surgically exposed lacrimal gland. Tear samples were subjected to protein quantification using the bicinchoninic acid (BCA) assay, and normalization prior to tryptic digestion and tandem mass tag (TMT) labeling. Quantitative proteomic analysis was performed by liquid chromatography–tandem mass spectrometry (LC–MS/MS). Differential protein expression analysis was conducted to identify significantly altered proteins. Selected candidate proteins along with their pathway-associated proteins were subsequently validated by Western blotting and immunofluorescence in a separate mouse cohort. Figure created with BioRender.com.

**Figure 1.**
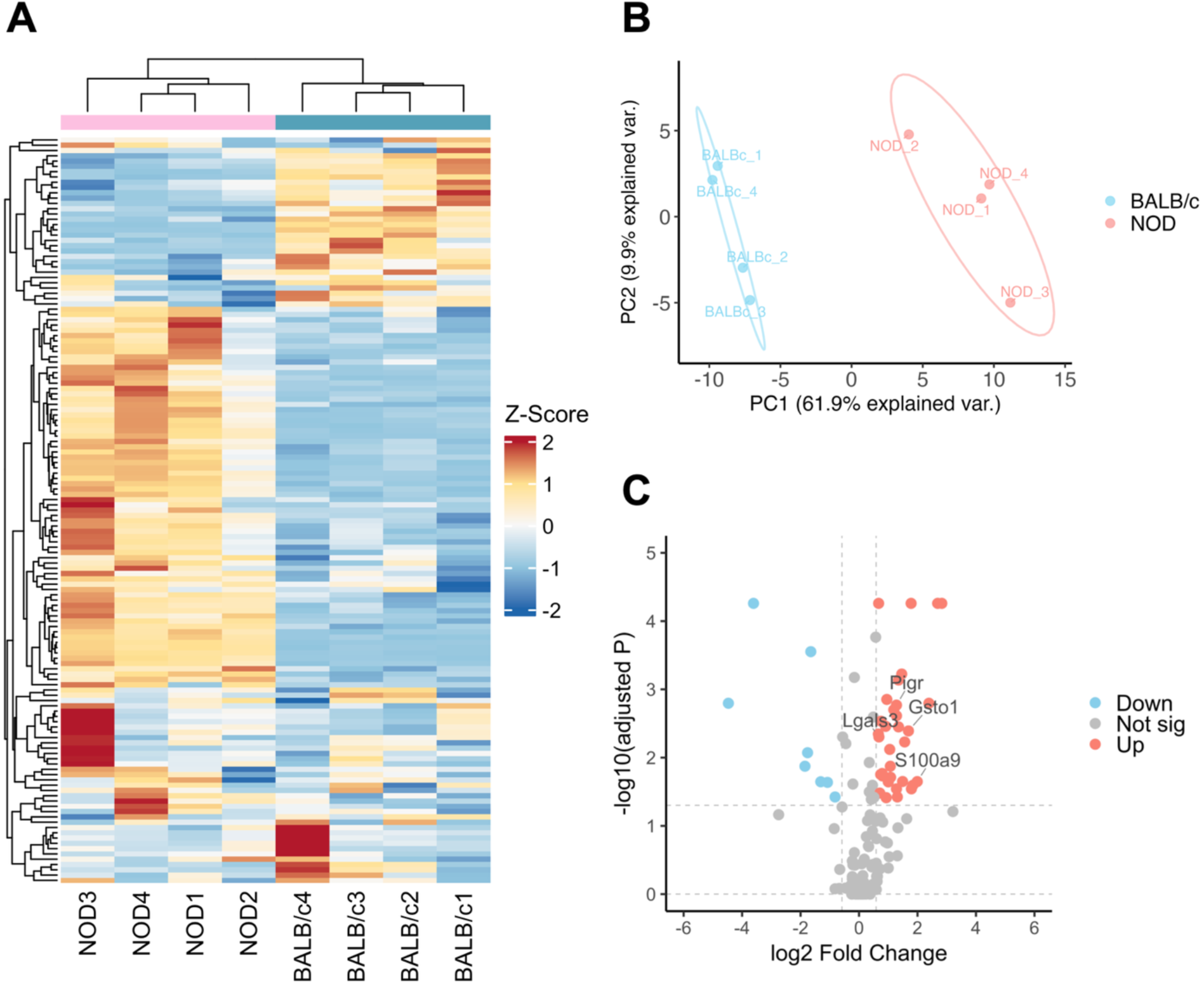
Proteome profiling of tears from male NOD and BALB/c mice. (**A**) Heatmap of detected tear proteins showing hierarchical clustering of NOD and BALB/c samples. Relative protein abundance values were z-score normalized across samples. (**B**) Principal component analysis (PCA) revealed a clear separation between NOD and BALB/c tear proteomes. (**C**) Volcano plot depicting differentially expressed proteins between NOD and BALB/c tears. Log2 fold change (NOD relative to BALB/c) is plotted against −log10(adjusted *P*). Proteins were considered significantly altered using adjusted *P* < 0.05 and absolute fold change > 1.5. Significantly upregulated and downregulated proteins are indicated, and selected upregulated candidates validated in subsequent experiments are labeled.

### Functional Enrichment Analysis indicates the involvement of DEPs in immune, inflammatory, and redox pathways

To gain functional insights into the DEPs in SjD-disease model NOD tears, GO enrichment analysis using Enrichr was conducted separately for upregulated and downregulated proteins across biological process (BP), molecular function (MF), and cellular component (CC) categories. Enriched GO terms such as mucosal immune response, RAGE receptor binding, secretory granule lumen, glutathione metabolic process, and regulation of cytokine production were primarily associated with proteins elevated in NOD mouse tears (**Fig. 2A**). These enrichments suggest that DEPs are associated with secretory compartments and/or involved in immune and inflammatory activation. Additionally, WikiPathways analysis also highlighted enrichment of proteins related to oxidative stress and redox regulation pathways, reinforcing findings from GO analysis and suggesting that both inflammatory and oxidative stress processes contribute to the altered NOD mouse tear proteome (**Fig. 2B**). GO analysis of downregulated proteins in SjD-disease model NOD mouse tears also highlighted pathways related to granule and vesicle lumina, suggesting changes in membrane trafficking pathways, while WikiPathways suggested downregulation of apoptosis modulation and oxidative stress response. Subsequent analyses focused primarily on the upregulated proteins, which showed stronger and more biologically consistent enrichment in pathways relevant to SjD-associated inflammation and redox dysregulation.

**Figure 2.**
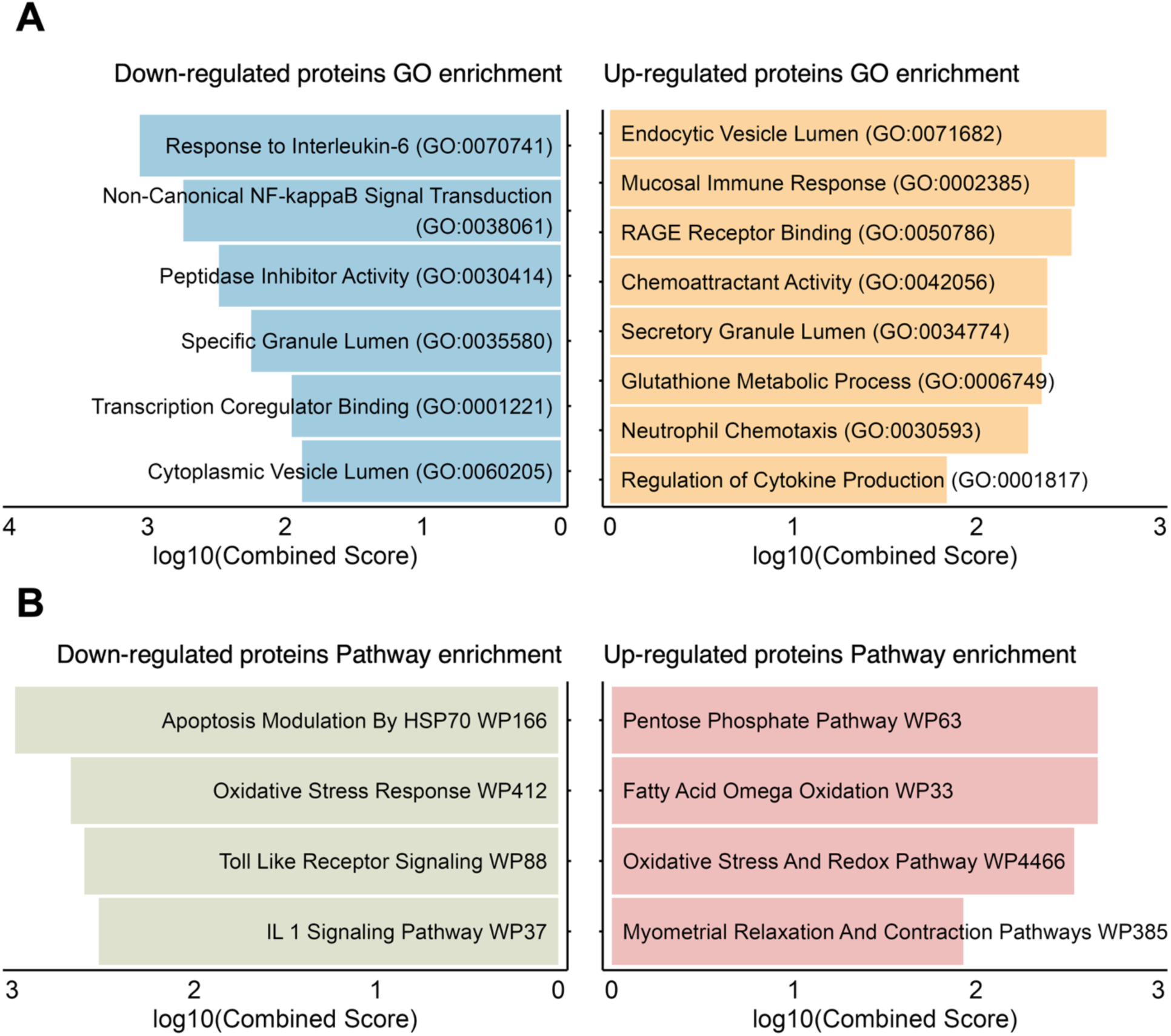
Functional enrichment analysis of differentially expressed tear proteins. (**A**) Gene Ontology (GO) enrichment analysis of proteins significantly up-regulated and down-regulated in male NOD mouse tears compared with male BALB/c mouse tears. (**B**) WikiPathways enrichment analysis of the same protein sets. Differentially expressed proteins were defined as previously stated. Enrichment analyses were performed using Enrichr, which calculates enrichment significance using a Fisher exact test relative to the respective background gene set. Bars represent log10(Combined Score). The Combined Score is defined by Enrichr as ln(*P*) × z-score, where *P* is the unadjusted enrichment *P* value and the z-score reflects deviation from the expected rank. Displayed terms met criteria of unadjusted *P* < 0.05 and Benjamini–Hochberg adjusted *P* < 0.1.

### Validation of selected tear proteomic hits in tears and LG

To further examine proteomic changes associated with the immune, inflammatory, and redox processes identified by enrichment analysis, four upregulated proteins including S100A9, glutathione S-transferase omega 1-1 (GSTO1-1), galectin-3 (Gal-3), and pIgR were selected for validation by Western blotting in tears from an independent cohort of mice. (**Fig. 3A and 3C**). S100A9 (also known as MRP14) is a member of the S100 calcium binding protein family and commonly exists as a heterodimer S100A8/A9 (also termed calprotectin) with S100A8^46,47^. S100A8/A9 is a potent damage-associated molecular pattern (DAMP) molecule implicated in inflammatory signaling and immune activation. S100A8/A9 levels were significantly increased in disease model NOD mouse tears compared to tears from healthy BALB/c mice (**Fig. 3A and 3C**). GSTO1-1 which functions in cellular detoxification and redox homeostasis by acting both as a thioltransferase and a reductase^48,49^ was also confirmed as significantly elevated in disease model NOD mouse tears relative to healthy BALB/c tears (**Fig. 3A and 3C**). Gal-3 is a β-galactoside-binding lectin involved in diverse inflammatory and immune responses. Gal-3 levels were also significantly increased in NOD mouse tears compared to BALB/c mouse tears (**Fig. 3A and 3C**). Finally, the secreted extracellular domain of pIgR, termed secretory component (SC), which is released into tears together with bound dimeric IgA (secretory IgA) upon its basolateral to apical transcytosis^15,16^, was likewise significantly increased in SjD disease model NOD mouse tears relative to healthy control tears (**Fig. 3A and 3C**). Many but not all proteins secreted into tears originate from the LG, reaching the tears through exocytosis or transcytosis from LGAC^13–16^. In the diseased LG in SjD, proteins present in tissue interstitium may also reach the tears due to the presence of leaky junctions between acinar and ductal epithelia associated with tissue inflammation and apoptosis^33^. The cornea, conjunctiva and meibomian glands also contribute components to the tear fluid^12,50^. To relate changes in the tear composition of SjD model NOD mice to disease processes occurring in the LG, we investigated whether the upregulated tear proteins of interest were also increased in the LG of the disease model mice. As shown in **Fig. 3B and 3D**, S100A8/A9, GSTO1-1, and Gal-3 were all significantly increased in LG lysates from NOD mice compared to BALB/c mice. Additionally, increased levels of both full-length pIgR and its cleaved extracellular domain, SC, were detected in NOD mouse LG lysates (**Fig. 3B and 3D**).

**Figure 3.**
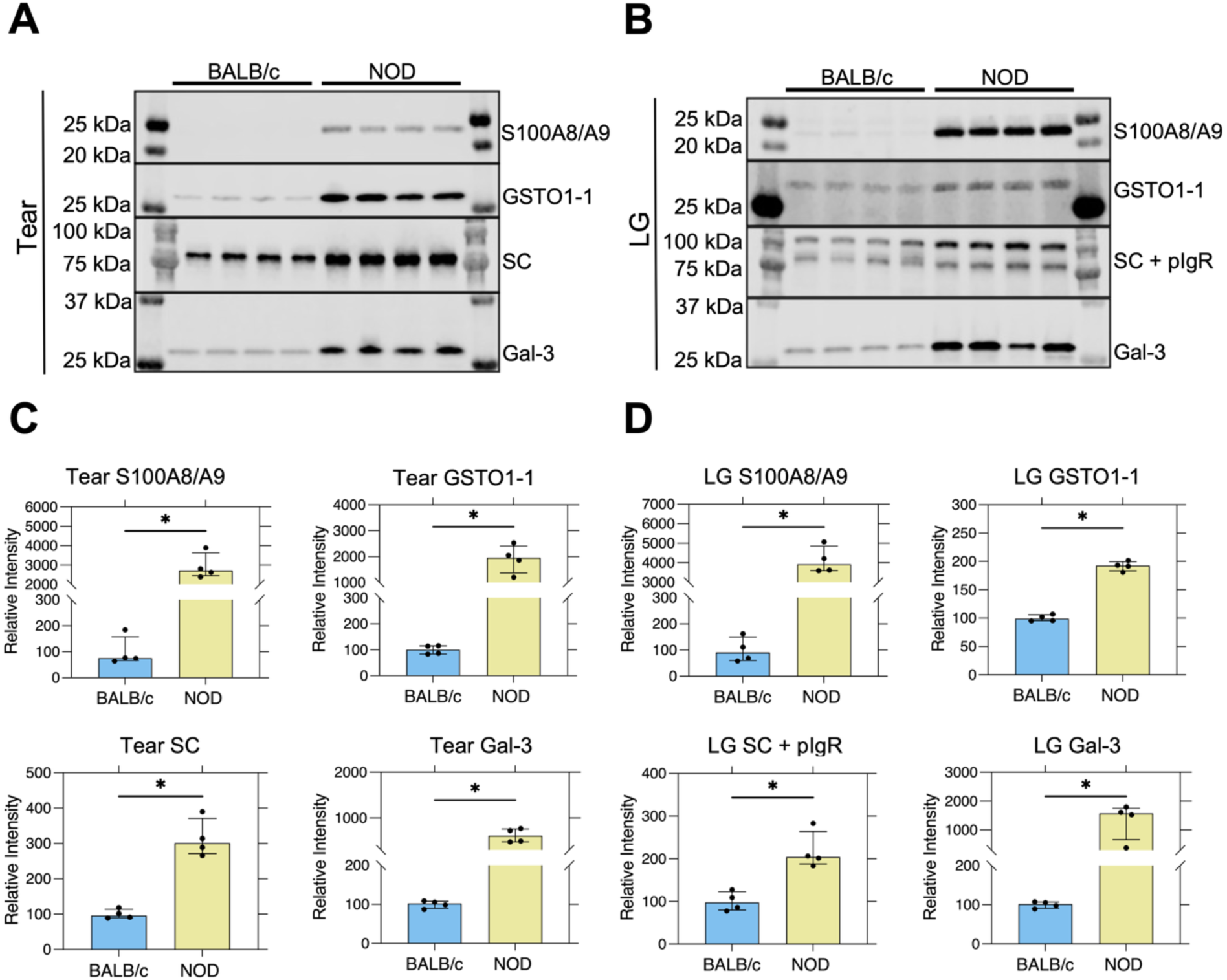
Western blotting validation of selected upregulated tear proteomic hits in tears and LG. (**A**), (**B**) Representative Western blots validating selected protein hits in tear samples (**A**) and LG lysates (**B**) from male BALB/c and NOD mice. Tear targets include S100A8/A9, GSTO1-1, free secretory component (SC), and galectin-3 (Gal-3). LG targets include S100A8/A9, GSTO1-1, SC and polymeric immunoglobulin receptor (pIgR), and Gal-3. Each lane represents one biological replicate. Tear samples were pooled from three mice per replicate. (**C**), (**D**) Quantification of Western blot signals for tear S100A8/A9, GSTO1-1, SC, and Gal-3; LG S100A8/A9, GSTO1-1, SC and pIgR, and Gal-3. Band intensities were normalized to total protein staining and are presented as relative intensity. Each dot represents one biological replicate (n = 4 per strain). Bars represent median ± IQR. Statistical comparisons were performed using the two-tailed Mann–Whitney U test. \**P* < 0.05.

To further examine the localization of these proteins within the LG, selected hits were also evaluated via immunofluorescence labeling of LG sections from 14-week-old male BALB/c and NOD mice (**Fig. 4**). In BALB/c mouse LG, S100A8/A9 labeling was minimal and limited to scattered myeloid-like cells. In NOD mouse LG, S100A8/A9-positive cells were more abundant within immune cell infiltrates. Gal-3 labeling also showed limited signal in BALB/c mouse LG, whereas intense labeling was observed in cells within inflammatory infiltrates in NOD mouse LG. In contrast, GSTO1-1 labeling was detected in a punctate pattern in LGAC in healthy BALB/c mouse LG, but this pattern was markedly altered in disease model NOD mouse LG, which showed GSTO1-1 labeling concentrated primarily beneath and within apical/lumenal acinar regions. This altered localization may reflect increased apical trafficking and secretion from LGAC, and may contribute to the elevated GSTO1-1 detected in NOD tears. pIgR immunofluorescence was largely detected at the basolateral membrane in BALB/c mouse LG acini, but the intensity of the signal was increased with a redistribution to the apical/lumenal regions in acini from NOD mouse LG, consistent with increased pIgR transcytosis.

**Figure 4.**
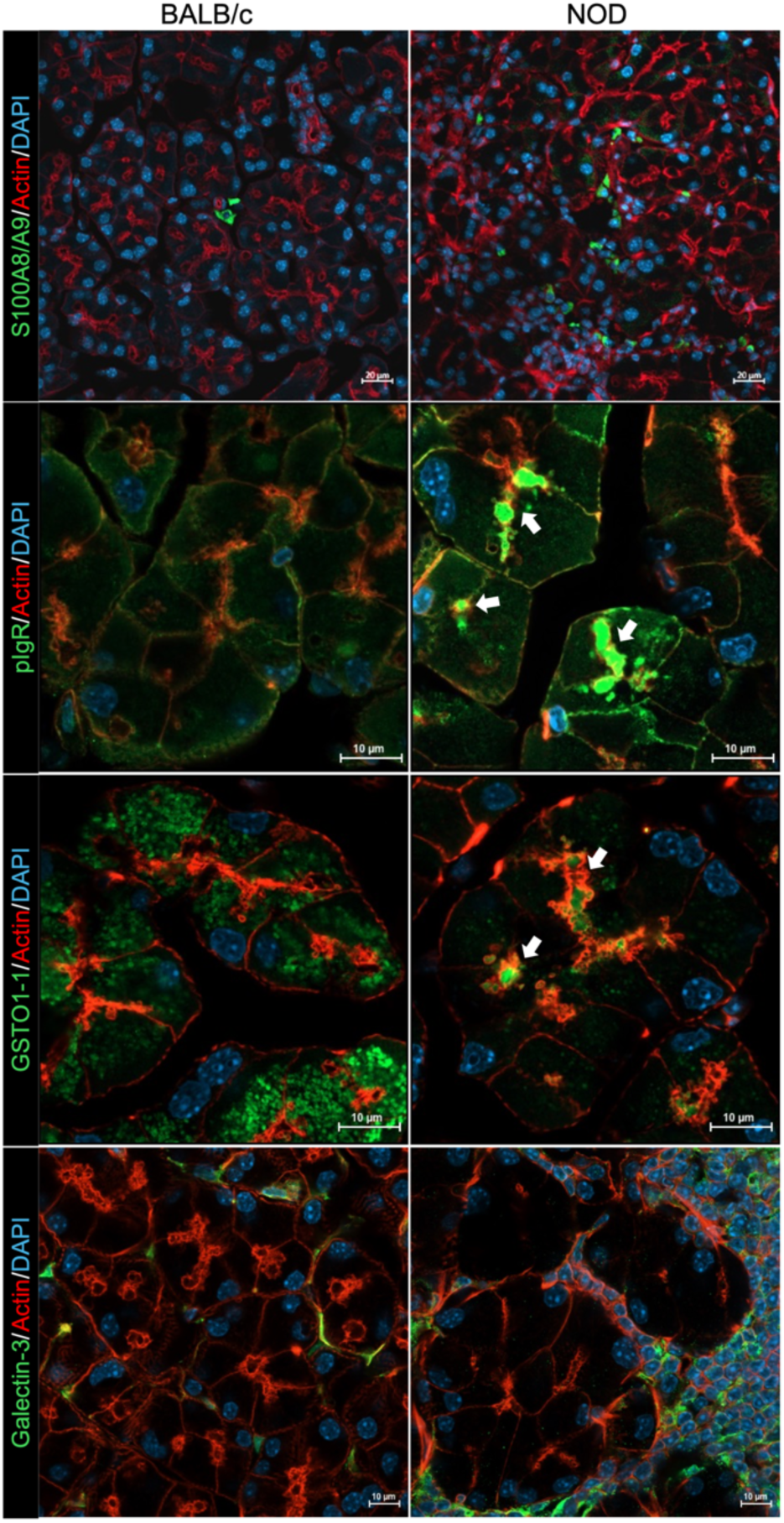
Immunofluorescence analysis of selected upregulated tear proteomic hits in male NOD mouse LG. Representative immunofluorescence images of male BALB/c and NOD mouse LG sections showing the localization of validated targets. Arrows indicate signal enrichment in apical/lumenal regions. Digital zoom varied among images to highlight differences in protein localization; scale bars are provided for reference. Scale bars: 20 μm for the top row and 10 μm for the remaining rows, as indicated.

### Increased S100A8/A9 is accompanied by elevated RAGE expression in NOD mouse LG

S100A8/A9 is implicated in pro-inflammatory signaling through RAGE (Receptor for Advanced Glycation End Products)^47^ (**Fig. 5A**). Given the increased expression level of S100A8/A9 in both NOD mouse tears and LG lysates, RAGE expression levels were evaluated in diseased NOD mouse and healthy BALB/c mouse LG. Western blotting demonstrated increased RAGE expression in NOD mouse LG compared to BALB/c mouse LG (**Fig. 5B**). Immunofluorescence labeling of the LG likewise showed minimal expression of RAGE in healthy BALB/c mouse LG, but higher expression in diseased NOD mouse LG, specifically in areas enriched in infiltrating lymphocytes (**Fig. 5C**).

**Figure 5.**
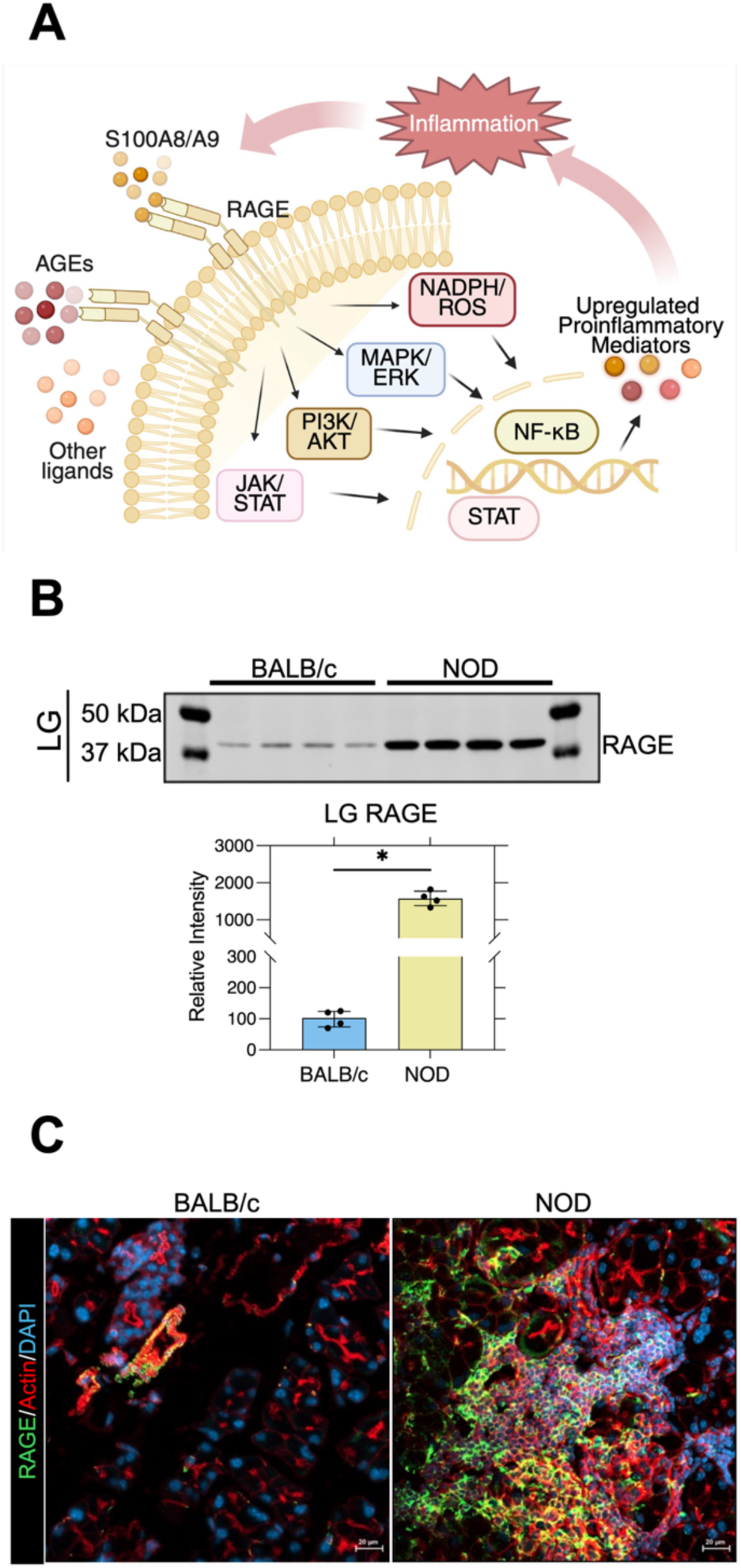
S100A8/A9–RAGE signaling is activated in NOD mouse LG. (**A**) Schematic illustration of the proposed S100A8/A9–RAGE signaling cascade in LG inflammation, highlighting downstream pathways and the amplification of inflammatory mediator expression. Figure created with BioRender.com. (**B**) Representative Western blots showing RAGE expression in LG lysates from male BALB/c and NOD mice, with quantification shown below. Each lane represents one biological replicate. Band intensities were normalized to total protein staining and are presented as relative intensity. Each dot represents one biological replicate (n = 4 per strain). Bars represent median ± IQR. Statistical comparisons were performed using the two-tailed Mann–Whitney U test. \**P* < 0.05. (**C**) Representative immunofluorescence images of LG sections showing RAGE (green) with actin (red) and DAPI (blue) from male BALB/c and NOD mice. Scale bars: 20 μm.

### Activation of NLRP3 inflammasome in NOD LG

GSTO1-1 has recently been implicated in the regulation of redox-imbalanced inflammatory pathways, including NLRP3 inflammasome activation^51,52^(**Fig. 6A**). Given the increased GSTO1-1 levels detected in both NOD mouse tears and LG lysates, we assessed the expression of other inflammasome activation-related proteins and downstream effectors in LG tissue. Western blotting showed increased NLRP3 protein abundance in NOD mouse LG compared to healthy BALB/c mouse LG (**Fig. 6B and 6C**). In addition, caspase-1, cleaved caspase-1, full-length gasdermin (GSDMD), cleaved N-terminal GSDMD, pro-IL-1β and cleaved IL-1β levels were all significantly increased in LG from NOD mice (**Fig. 6B and 6C**). RT-qPCR analysis further confirmed increased gene expression of *Il1b*, *Il18*, and *Il18r*, cytokine mediators of NLRP3 activation, in NOD mouse LG relative to healthy BALB/c controls (**Fig. 6D**).

**Figure 6.**
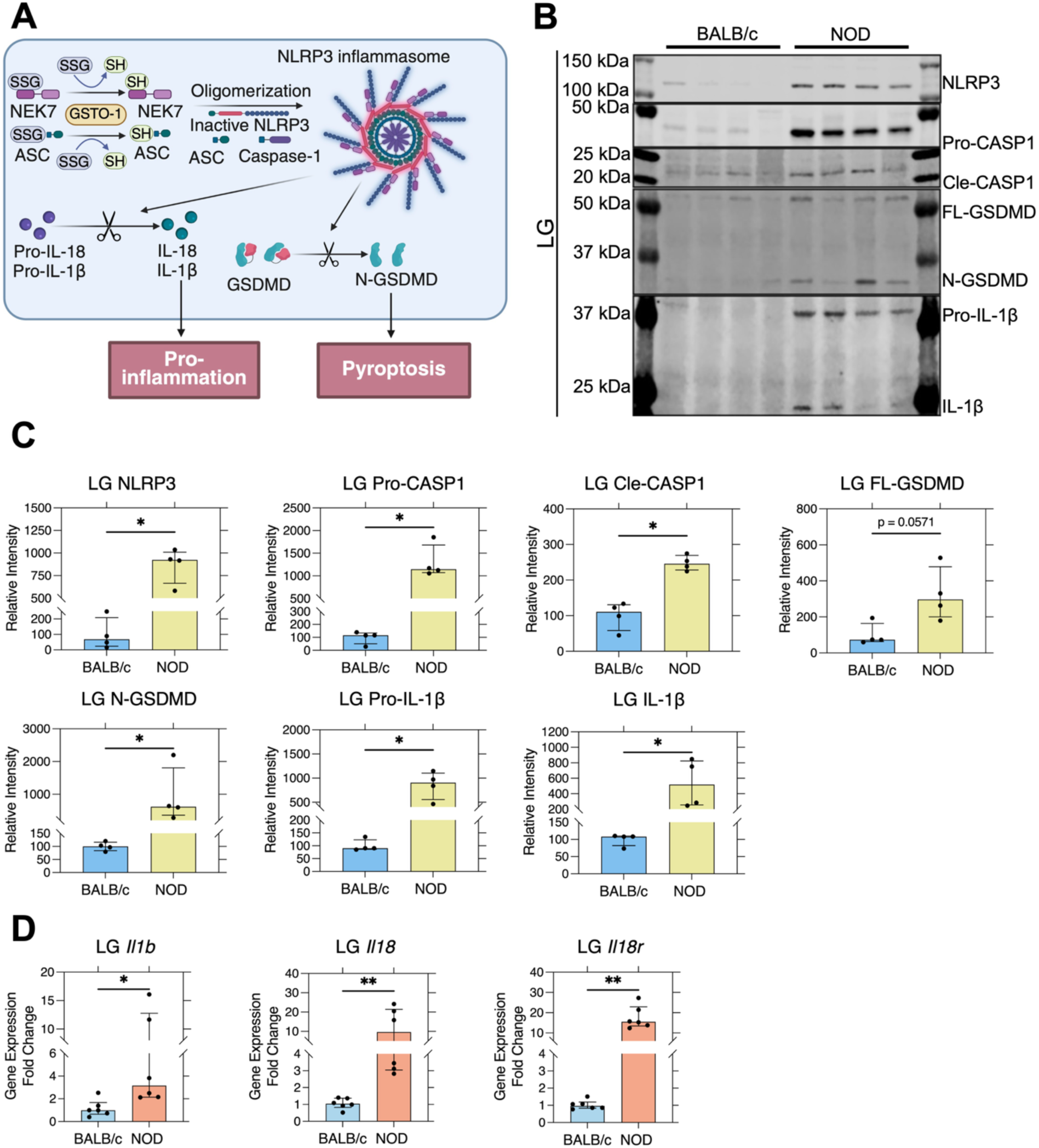
NLRP3 inflammasome activation in male NOD mouse LG. (**A**) Schematic illustration of the NLRP3 inflammasome activation leading to pyroptosis. Figure created with BioRender.com. (**B**) Representative Western blots of LG lysates from male BALB/c and NOD mice showing NLRP3, pro-caspase-1, cleaved caspase-1, full-length GSDMD (FL-GSDMD), N-terminal GSDMD (N-GSDMD), pro-IL-1β, and IL-1β. Each lane represents one biological replicate. (**C**) Quantification of Western blots signals. Band intensities were normalized to total protein staining and are presented as relative intensity. Each dot represents one biological replicate (n = 4 per strain). Bars represent median ± IQR. Group comparisons were performed using the two-tailed Mann–Whitney U test. \**P* < 0.05. (**D**) Relative mRNA expression of *Il1b*, *Il18*, and *Il18r* in LG. Each dot represents one biological replicate from an individual mouse (n = 6 per strain). Bars represent median ± IQR. Statistical comparisons were performed using the two-tailed Mann–Whitney U test. \**P* < 0.05, \*\**P* < 0.01.

## DISCUSSION

Quantitative tear proteomics revealed broad tear proteome alterations in the male NOD mouse model of SjD-like autoimmune dacryoadenitis. Identified tear DEPs were enriched for immune and inflammatory responses and redox-associated biological processes and pathways, consistent with established features of SjD-associated autoimmune dacryoadenitis, in which lymphocytic infiltration and chronic inflammation impair normal LG secretory functions and alter tear production^3,18,19,31–37^. Selected tear DEPs were also validated as increased in diseased LG, indicating that the inflamed LG is the major contributor to these tear proteome alterations. This reinforces the premise that tears can provide a noninvasive readout of pathological processes within the LG and can be leveraged for biomarker discovery in SjD. A recent proteomic study of chronically inflamed LG from NOR/LtJ mice, a NOD-derived strain, also identified broad disease-associated proteome alterations, further supporting the extensive molecular shift in SjD-associated autoimmune dacryoadenitis^53^.

Male NOD mice are commonly used in studies of ocular symptoms of SjD, as they spontaneously develop autoimmune dacryoadenitis and tear hyposecretion characteristic of human disease^18,31–37^. In these mice, detectable immune cell infiltration forms in LG as early as 8-10 weeks of age. By 14 weeks of age, male NOD mice exhibit intermediate glandular inflammation and functional changes that are distinct from the end stage, severe disease seen in mice at older ages^31,35,54^. Thus, the 14-week-old male NOD mice used here likely reflect early-to-intermediate level SjD in the LG, making this model well suited for identification of tear biomarkers and disease-associated pathways that initiate and promote SjD at disease onset, when therapeutic interventions may improve glandular inflammation and function^34^. S100A8/A9 has been widely studied in autoimmune diseases, including rheumatoid arthritis (RA), systemic lupus erythematosus (SLE), and SjD, where it is associated with inflammatory responses and disease progression^47,55^. In SjD, prior studies have reported markedly elevated S100A8/A9 in patient PBMCs, serum, and parotid saliva^56–58^, and its expression has also been linked to disease severity and reduced tear production^56,57^. As danger signals released during cell stress, cell injury, or immune cell activation, DAMPs perpetuate local inflammatory signaling and contribute to persistent glandular inflammation^21,59^. Multiple DAMPs, including HMGB1, IL-33, IL-1α, ATP, histones, and S100A8/A9, have been implicated in SjD through studies of patient serum, tears, saliva, PBMCs, and glandular tissues, as well as experimental mouse models^21,56–58^. S100A8/A9 induced in the LG under conditions of chronic inflammation may be released into the extracellular space, amplifying inflammatory responses through pattern-recognition receptors (PRR) such as RAGE^56,60,61^. RAGE provides a receptor-mediated mechanism through which extracellular DAMPs may sustain and amplify local inflammatory signaling^60^. Prior studies in SjD have suggested increased RAGE expression in glandular tissue, with strong labeling detected in SjD patient SG epithelial and myoepithelial cells^62^. Importantly, RAGE signaling is associated not only with recruitment of immune cells to infiltrates and persistence of infiltrates, but also to activation of downstream MAPK-and NF-κB-related signaling pathways, thus promoting pro-inflammatory cytokine production, oxidative stress, and sustained inflammatory responses^60^. In this study, the marked increase in S100A8/A9 and RAGE enrichment in NOD mouse LG in immune infiltrates is consistent with the contribution of this signaling pathway to chronic inflammation within the diseased gland. Thus, S100A8/A9 may serve not only as a biomarker of autoimmune inflammation of the LG but may actively participate in disease propagation. S100A6, another S100 family of calcium-binding proteins, was also significantly elevated in NOD mouse tears in the proteomics analysis, and has previously been implicated in cellular stress responses and inflammatory signaling^63,64^. Its increased abundance together with elevated S100A8/A9 and RAGE is consistent with the presence of a broader DAMP-associated inflammatory environment within the diseased LG.

Another notable finding was the increased abundance of Gal-3 in NOD mouse tears and LG. Gal-3 is involved in inflammatory and immune responses and has previously been implicated in SjD^65^. Elevated Gal-3 has been reported in serum from patients with SjD and is correlated with markers of systemic inflammation including C-reactive protein (CRP), fibrinogen, IgG, and IL-17^65^. In addition, increased tear Gal-3 has been reported in DED associated with epithelial dysfunction and tear film instability^66^. Consistent with these observations, Gal-3 immunofluorescence in this study was primarily localized in lymphocytic infiltrates within NOD mouse LG, suggesting that infiltrating immune cells may contribute to its increased abundance in diseased tissue. Also related to inflammation, proteomics analysis identified increased beta-2 microglobulin (B2M) in NOD mouse tears. B2M is a component of the major histocompatibility complex class I (MHC-I) molecule. It is elevated in serum and saliva from patients with SjD^67,68^, and is also associated with disease activity in SjD^68,69^.

In addition to identification of tear proteins linked to inflammation in the diseased NOD mouse LG as noted above, oxidative stress contributes to development of SjD, with recent work showing altered antioxidant defenses in patients with SjD^29^. In this context, GSTO1-1 was markedly increased in both tears and LG lysates from diseased mice and may represent a tear biomarker of particular interest. GSTO1-1 is an omega-class glutathione transferase with reductase and thioltransferase activity, and its related redox reactions have been implicated in modulation of intracellular signaling pathways, including NF-κB-associated pathways^70,71^. In recent years, increasing evidence has suggested a pro-inflammatory role of GSTO1-1 in different disease models^72^. In SjD, a recent multi-omics analysis identified increased GSTO1-1 gene expression in patient samples, suggesting its potential role as a diagnostic biomarker for SjD^73^. In our study, GSTO1-1 showed a punctate pattern in healthy BALB/c LG tissue, whereas the signal in NOD mouse LG appeared to be enriched within the apical/lumenal regions of acini. This shift may suggest that the acinar cells not only upregulate the expression of GSTO1-1 but also redistribute this enzyme to their apical/lumenal membrane compartments for active secretion under conditions of stress or inflammation. Consistent with this premise, GSTO1-1 is detectable in extracellular biofluids, including sputum supernatants, bronchoalveolar lavage fluid, and plasma, where it has been proposed to contribute to extracellular redox balance^74,75^, reflecting a biologically meaningful release rather than simply leakage from damaged cells^74,76^. The observed increase of GSTO1-1 in NOD mouse tears may partially reflect a redox homeostasis-associated redistribution and increased secretion in response to chronic inflammatory stress.

Although GSTO1-1 may serve a partially protective role in redox homeostasis in biofluids, it is also a known activator of the NLRP3 inflammasome through deglutathionylating of key inflammasome components including apoptosis-associated speck-like protein containing a CARD (ASC) and NIMA related kinase 7 (NEK7)^51,52^. The NLRP3 inflammasome functions as an intracellular PRR that senses endogenous danger signals, and once activated, drives the activation of caspase-1, maturation of IL-1β and IL-18 and cleavage of GSDMD^28^. Such activation, when sustained, can trigger pyroptosis, a lytic inflammatory form of regulated cell death characterized by membrane pore formation, cell swelling and release of proinflammatory cytokines^28^. All these components of NLRP3 inflammasome activation are detected in the NOD mouse LG in our study. Our findings are consistent with prior studies in dry eye and SjD which have reported increased inflammasome components in patient tears and conjunctival epithelial cells^27^, while studies in SG in SjD have also detected elevated NLRP3 inflammasome in the labial gland and SG of SjD patients and the SG of female NOD mice^77,78^. In parallel, studies in LG epithelial cells have shown that both acute and chronic inflammation can induce sustained NLRP3 inflammasome activation and pyroptosis in association with epithelial dysfunction and disease progression in SjD-like mouse models^30^. Our findings provide additional evidence for NLRP3 inflammasome-driven pyroptotic cell death in the NOD disease model, and to our knowledge, provides the first evidence that tear GSTO1-1 may reflect LG NLRP3 inflammasome activation and pyroptosis-associated epithelial dysfunction in the LG in SjD.

pIgR and SC were increased in both NOD mouse tears and LG, and showed a detectable accumulation in the apical/lumenal regions of acinar cells within the diseased LG. Under physiological conditions, pIgR mediates transcytosis of dimeric IgA and IgM from the basolateral to apical surfaces of acinar cells where it is cleaved. The released extracellular domain of pIgR bound to dimeric IgA is released as free SC into tear fluid, and constitutes a major component of mucosal immunity^15,16^. pIgR expression is known to be upregulated by pro-inflammatory cytokines implicated in SjD, including IL-1, IL-17, IFN-γ, and TNF-α through NF-κB-and IRF-1-associated pathways^79^, which may partially account for its increased levels in tears and tissue in disease. We have recently demonstrated the formation of ectopic lymphoid structures (ELS) in the LG of male NOD mice, which are capable of sustaining local autoantibody production^37,80^. We have further shown that IgA autoantibodies are selectively increased in tears, relative to their levels in serum, and are presumably produced at least in part by LG ELS^37^. The increased abundance of dimeric IgA autoantibodies in NOD mouse LG may further drive increased pIgR expression and transcytosis leading to increased secretion of SC in disease.

Some limitations should be considered regarding the current study. Although tears provide a readily accessible and minimally invasive biofluid, the tear proteome reflects not only LG secretory functions but also contributions from other ocular surface epithelium^50^. As is common in discovery proteomics, the sample cohort used in the present study was modest in sample size, which may reduce our ability to detect more subtle changes and mask variability present in the disease mouse tear proteome. Moreover, the models used in this study were evaluated at a fixed time point and may not have captured important temporal changes in tear composition that might reflect specific disease stages of the LG. Finally, while the relationship of tear proteome changes to the occurrence of pathological processes in the LG was demonstrated here for tear S100A8/A9 and LG RAGE activation and increased tear GSTO1-1 and LG NLRP3 inflammasome activation, the overall contribution of these pathways to disease pathogenesis is still unclear.

In conclusion, the present study shows that tear proteome remodeling in the NOD model of ocular SjD reflects biologically meaningful changes occurring within the diseased LG. Increased tear S100A8/A9, GSTO1-1, and pIgR, together with validation that their downstream pathways are altered in the LG, indicate that tear proteins can capture distinct features of SjD-associated LG pathology, including inflammatory propagation, redox-associated stress responses, and altered epithelial cell transcytosis. Meanwhile, the demonstration of NLRP3 inflammasome activation and pyroptotic cell death signaling in diseased LG suggests that inflammasome-driven epithelial dysfunction may also contribute to disease development and/or progression. These findings support tear proteomics as a useful approach for biomarker discovery in SjD and raise the possibility that selected tear proteins, particularly GSTO1-1, may serve as informative readouts of active pathological processes within the LG in patients with SjD.

## ACKNOWLEDGEMENTS

This work was supported by NEI R01 grants EY011386 and EY026635 to SHA, and an NEI P30EY029220 to the Department of Ophthalmology at the USC Keck School of Medicine, and an Unrestricted Grant from Research to Prevent Blindness to the Department of Ophthalmology at the USC Keck School of Medicine. Also supported by NIDDK F31 grant F31DK138790-01 to MA.

## DATA AVAILABILITY

The mass spectrometry proteomics data have been deposited to the ProteomeXchange Consortium via the PRIDE partner repository (https://www.ebi.ac.uk/pride) with the dataset identifier PXD081121.

## Commercial Relationships Disclosure

None

## SUPPLEMENTARY MATERIAL

**Supplementary Table S1.** Antibodies used for validation experiments.

## REFERENCES

1. Brito-Zeron P, Baldini C, Bootsma H, et al. Sjogren syndrome. Nat Rev Dis Primers. Jul 7 2016;2:16047. doi:10.1038/nrdp.2016.47

2. Akpek EK, Bunya VY, Saldanha IJ. Sjogren’s Syndrome: More Than Just Dry Eye. Cornea. May 2019;38(5):658–661. doi:10.1097/ICO.0000000000001865

3. Nguyen CQ, Peck AB. Unraveling the pathophysiology of Sjogren syndrome-associated dry eye disease. Ocul Surf. Jan 2009;7(1):11–27. doi:10.1016/s1542-0124(12)70289-6

4. Shiboski CH, Shiboski SC, Seror R, et al. 2016 American College of Rheumatology/European League Against Rheumatism Classification Criteria for Primary Sjogren’s Syndrome: A Consensus and Data-Driven Methodology Involving Three International Patient Cohorts. Arthritis Rheumatol. Jan 2017;69(1):35–45. doi:10.1002/art.39859

5. Huang YT, Lu TH, Chou PL, Weng MY. Diagnostic Delay in Patients with Primary Sjogren’s Syndrome: A Population-Based Cohort Study in Taiwan. Healthcare (Basel). Mar 23 2021;9(3)doi:10.3390/healthcare9030363

6. Ritter J, Chen Y, Stefanski AL, Dorner T. Current and future treatment in primary Sjogren’s syndrome - A still challenging development. Joint Bone Spine. Nov 2022;89(6):105406. doi:10.1016/j.jbspin.2022.105406

7. van Nimwegen JF, Moerman RV, Sillevis Smitt N, Brouwer E, Bootsma H, Vissink A. Safety of treatments for primary Sjogren’s syndrome. Expert Opin Drug Saf. 2016;15(4):513–524. doi:10.1517/14740338.2016.1146676

8. Verstappen GM, Pringle S, Bootsma H, Kroese FGM. Epithelial-immune cell interplay in primary Sjogren syndrome salivary gland pathogenesis. Nat Rev Rheumatol. Jun 2021;17(6):333–348. doi:10.1038/s41584-021-00605-2

9. Soret P, Le Dantec C, Desvaux E, et al. A new molecular classification to drive precision treatment strategies in primary Sjogren’s syndrome. Nat Commun. Jun 10 2021;12(1):3523. doi:10.1038/s41467-021-23472-7

10. Chaly Y, Barr JY, Sullivan DA, Thomas HE, Brodnicki TC, Lieberman SM. Type I Interferon Signaling Is Required for Dacryoadenitis in the Nonobese Diabetic Mouse Model of Sjogren Syndrome. Int J Mol Sci. Oct 20 2018;19(10)doi:10.3390/ijms19103259

11. McDermott AM. Antimicrobial compounds in tears. Exp Eye Res. Dec 2013;117:53–61. doi:10.1016/j.exer.2013.07.014

12. Willcox MDP, Argueso P, Georgiev GA, et al. TFOS DEWS II Tear Film Report. Ocul Surf. Jul 2017;15(3):366–403. doi:10.1016/j.jtos.2017.03.006

13. Wu K, Jerdeva GV, da Costa SR, Sou E, Schechter JE, Hamm-Alvarez SF. Molecular mechanisms of lacrimal acinar secretory vesicle exocytosis. Exp Eye Res. Jul 2006;83(1):84–96. doi:10.1016/j.exer.2005.11.009

14. Dartt DA. Neural regulation of lacrimal gland secretory processes: relevance in dry eye diseases. Prog Retin Eye Res. May 2009;28(3):155–177. doi:10.1016/j.preteyeres.2009.04.003

15. Knop E, Knop N, Claus P. Local production of secretory IgA in the eye-associated lymphoid tissue (EALT) of the normal human ocular surface. Invest Ophthalmol Vis Sci. Jun 2008;49(6):2322–2329. doi:10.1167/iovs.07-0691

16. Xu S, Ma L, Evans E, Okamoto CT, Hamm-Alvarez SF. Polymeric immunoglobulin receptor traffics through two distinct apically targeted pathways in primary lacrimal gland acinar cells. J Cell Sci. Jun 15 2013;126(Pt 12):2704–2717. doi:10.1242/jcs.122242

17. Conrady CD, Joos ZP, Patel BC. Review: The Lacrimal Gland and Its Role in Dry Eye. J Ophthalmol. 2016;2016:7542929. doi:10.1155/2016/7542929

18. Fu R, Guo H, Janga S, et al. Cathepsin S activation contributes to elevated CX3CL1 (fractalkine) levels in tears of a Sjogren’s syndrome murine model. Sci Rep. Jan 29 2020;10(1):1455. doi:10.1038/s41598-020-58337-4

19. Edman MC, Janga SR, Meng Z, et al. Increased Cathepsin S activity associated with decreased protease inhibitory capacity contributes to altered tear proteins in Sjogren’s Syndrome patients. Sci Rep. Jul 23 2018;8(1):11044. doi:10.1038/s41598-018-29411-9

20. Nakamura H, Tanaka T, Pranzatelli T, et al. Lysosome-associated membrane protein 3 misexpression in salivary glands induces a Sjogren’s syndrome-like phenotype in mice. Ann Rheum Dis. Aug 2021;80(8):1031–1039. doi:10.1136/annrheumdis-2020-219649

21. Ming B, Zhu Y, Zhong J, Dong L. Immunopathogenesis of Sjogren’s syndrome: Current state of DAMPs. Semin Arthritis Rheum. Oct 2022;56:152062. doi:10.1016/j.semarthrit.2022.152062

22. Fulton DJR, Li X, Bordan Z, et al. Galectin-3: A Harbinger of Reactive Oxygen Species, Fibrosis, and Inflammation in Pulmonary Arterial Hypertension. Antioxid Redox Signal. Nov 10 2019;31(14):1053–1069. doi:10.1089/ars.2019.7753

23. Riviere E, Chivasso C, Pascaud J, et al. Hyperosmolar environment and salivary gland epithelial cells increase extra-cellular matrix remodeling and lymphocytic infiltration in Sjogren’s syndrome. Clin Exp Immunol. Apr 7 2023;212(1):39–51. doi:10.1093/cei/uxad020

24. Rizzo C, Grasso G, Destro Castaniti GM, Ciccia F, Guggino G. Primary Sjogren Syndrome: Focus on Innate Immune Cells and Inflammation. Vaccines (Basel). Jun 3 2020;8(2)doi:10.3390/vaccines8020272

25. Parisis D, Chivasso C, Perret J, Soyfoo MS, Delporte C. Current State of Knowledge on Primary Sjogren’s Syndrome, an Autoimmune Exocrinopathy. J Clin Med. Jul 20 2020;9(7)doi:10.3390/jcm9072299

26. Zhuang D, Misra SL, Mugisho OO, Rupenthal ID, Craig JP. NLRP3 Inflammasome as a Potential Therapeutic Target in Dry Eye Disease. Int J Mol Sci. Jun 29 2023;24(13)doi:10.3390/ijms241310866

27. Niu L, Zhang S, Wu J, Chen L, Wang Y. Upregulation of NLRP3 Inflammasome in the Tears and Ocular Surface of Dry Eye Patients. PLoS One. 2015;10(5):e0126277. doi:10.1371/journal.pone.0126277

28. Carnazzo V, Rigante D, Restante G, Basile V, Pocino K, Basile U. The entrenchment of NLRP3 inflammasomes in autoimmune disease-related inflammation. Autoimmun Rev. Jun 24 2025;24(7):103815. doi:10.1016/j.autrev.2025.103815

29. Benchabane S, Sour S, Zidi S, et al. Exploring the relationship between oxidative stress status and inflammatory markers during primary Sjogren’s syndrome: A new approach for patient monitoring. Int J Immunopathol Pharmacol. Jan-Dec 2024;38:3946320241263034. doi:10.1177/03946320241263034

30. Delcroix V, Mauduit O, Yang M, et al. Lacrimal Gland Epithelial Cells Shape Immune Responses through the Modulation of Inflammasomes and Lipid Metabolism. Int J Mol Sci. Feb 21 2023;24(5)doi:10.3390/ijms24054309

31. Doyle ME, Boggs L, Attia R, et al. Autoimmune dacryoadenitis of NOD/LtJ mice and its subsequent effects on tear protein composition. Am J Pathol. Oct 2007;171(4):1224–1236. doi:10.2353/ajpath.2007.070388

32. Schenke-Layland K, Xie J, Angelis E, et al. Increased degradation of extracellular matrix structures of lacrimal glands implicated in the pathogenesis of Sjogren’s syndrome. Matrix Biol. Jan 2008;27(1):53–66. doi:10.1016/j.matbio.2007.07.005

33. Umazume T, Thomas WM, Campbell S, et al. Lacrimal Gland Inflammation Deregulates Extracellular Matrix Remodeling and Alters Molecular Signature of Epithelial Stem/Progenitor Cells. Invest Ophthalmol Vis Sci. Dec 2015;56(13):8392–8402. doi:10.1167/iovs.15-17477

34. Klinngam W, Janga SR, Lee C, et al. Inhibition of Cathepsin S Reduces Lacrimal Gland Inflammation and Increases Tear Flow in a Mouse Model of Sjogren’s Syndrome. Sci Rep. Jul 2 2019;9(1):9559. doi:10.1038/s41598-019-45966-7

35. Kakan SS, Edman MC, Yao A, et al. Tear miRNAs Identified in a Murine Model of Sjogren’s Syndrome as Potential Diagnostic Biomarkers and Indicators of Disease Mechanism. Front Immunol. 2022;13:833254. doi:10.3389/fimmu.2022.833254

36. Singh Kakan S, Li X, Edman MC, Okamoto CT, Hjelm BE, Hamm-Alvarez SF. The miRNA Landscape of Lacrimal Glands in a Murine Model of Autoimmune Dacryoadenitis. Invest Ophthalmol Vis Sci. Apr 3 2023;64(4):1. doi:10.1167/iovs.64.4.1

37. Singh Kakan S, Abdelhamid S, Ju Y, et al. Serum and tear autoantibodies from NOD and NOR mice as potential diagnostic indicators of local and systemic inflammation in Sjogren’s disease. Front Immunol. 2024;15:1516330. doi:10.3389/fimmu.2024.1516330

38. Lin Y, Zhang Y, Shi K, Wu H, Ou S. Advances in clinical examination of lacrimal gland. Front Med (Lausanne). 2023;10:1257209. doi:10.3389/fmed.2023.1257209

39. Li B, Sheng M, Li J, et al. Tear proteomic analysis of Sjogren syndrome patients with dry eye syndrome by two-dimensional-nano-liquid chromatography coupled with tandem mass spectrometry. Sci Rep. Aug 27 2014;4:5772. doi:10.1038/srep05772

40. Kuleshov MV, Jones MR, Rouillard AD, et al. Enrichr: a comprehensive gene set enrichment analysis web server 2016 update. Nucleic Acids Res. Jul 8 2016;44(W1):W90–97. doi:10.1093/nar/gkw377

41. Chen EY, Tan CM, Kou Y, et al. Enrichr: interactive and collaborative HTML5 gene list enrichment analysis tool. BMC Bioinformatics. Apr 15 2013;14:128. doi:10.1186/1471-2105-14-128

42. Gene Ontology C, Aleksander SA, Balhoff J, et al. The Gene Ontology knowledgebase in 2023. Genetics. May 4 2023;224(1)doi:10.1093/genetics/iyad031

43. Martens M, Ammar A, Riutta A, et al. WikiPathways: connecting communities. Nucleic Acids Res. Jan 8 2021;49(D1):D613–D621. doi:10.1093/nar/gkaa1024

44. Xie Z, Bailey A, Kuleshov MV, et al. Gene Set Knowledge Discovery with Enrichr. Curr Protoc. Mar 2021;1(3):e90. doi:10.1002/cpz1.90

45. Bankhead P, Loughrey MB, Fernandez JA, et al. QuPath: Open source software for digital pathology image analysis. Sci Rep. Dec 4 2017;7(1):16878. doi:10.1038/s41598-017-17204-5

46. Rammes A, Roth J, Goebeler M, Klempt M, Hartmann M, Sorg C. Myeloid-related protein (MRP) 8 and MRP14, calcium-binding proteins of the S100 family, are secreted by activated monocytes via a novel, tubulin-dependent pathway. J Biol Chem. Apr 4 1997;272(14):9496–9502. doi:10.1074/jbc.272.14.9496

47. Wang X, Luo Y, Zhou Q, Ma J. The roles of S100A8/A9 and S100A12 in autoimmune diseases: Mechanisms, biomarkers, and therapeutic potential. Autoimmun Rev. Dec 18 2025;24(12):103920. doi:10.1016/j.autrev.2025.103920

48. Board PG, Coggan M, Chelvanayagam G, et al. Identification, characterization, and crystal structure of the Omega class glutathione transferases. J Biol Chem. Aug 11 2000;275(32):24798–24806. doi:10.1074/jbc.M001706200

49. Schmuck EM, Board PG, Whitbread AK, et al. Characterization of the monomethylarsonate reductase and dehydroascorbate reductase activities of Omega class glutathione transferase variants: implications for arsenic metabolism and the age-at-onset of Alzheimer’s and Parkinson’s diseases. Pharmacogenet Genomics. Jul 2005;15(7):493–501. doi:10.1097/01.fpc.000016572581559.e3

50. Jumblatt MM, Imbert Y, Young WW, Jr., Foulks GN, Steele PS, Demuth DR. Glycoprotein 340 in normal human ocular surface tissues and tear film. Infect Immun. Jul 2006;74(7):4058–4063. doi:10.1128/IAI.01951-05

51. Li S, Wang L, Xu Z, et al. ASC deglutathionylation is a checkpoint for NLRP3 inflammasome activation. J Exp Med. Sep 6 2021;218(9)doi:10.1084/jem.20202637

52. Hughes MM, Hooftman A, Angiari S, et al. Glutathione Transferase Omega-1 Regulates NLRP3 Inflammasome Activation through NEK7 Deglutathionylation. Cell Rep. Oct 1 2019;29(1):151–161 e155. doi:10.1016/j.celrep.2019.08.072

53. Toribio D, Morokuma J, Pellino D, Hardt M, Zoukhri D. Quantitative Changes in the Proteome of Chronically Inflamed Lacrimal Glands From a Sjogren’s Disease Animal Model. Invest Ophthalmol Vis Sci. Apr 1 2025;66(4):44. doi:10.1167/iovs.66.4.44

54. Ohno Y, Satoh K, Shitara A, Into T, Kashimata M. Arginase 1 is involved in lacrimal hyposecretion in male NOD mice, a model of Sjogren’s syndrome, regardless of dacryoadenitis status. J Physiol. Nov 2020;598(21):4907–4925. doi:10.1113/JP280090

55. Kopec-Medrek M, Widuchowska M, Kucharz EJ. Calprotectin in rheumatic diseases: a review. Reumatologia. 2016;54(6):306–309. doi:10.5114/reum.2016.64907

56. Wei Y, Sun M, Zhang X, et al. S100A8/A9 Promotes Dendritic Cell-Mediated Th17 Cell Response in Sjogren’s Dry Eye Disease by Regulating the Acod1/STAT3 Pathway. Invest Ophthalmol Vis Sci. Jan 2 2025;66(1):35. doi:10.1167/iovs.66.1.35

57. Nicaise C, Weichselbaum L, Schandene L, et al. Phagocyte-specific S100A8/A9 is upregulated in primary Sjogren’s syndrome and triggers the secretion of pro-inflammatory cytokines in vitro. Clin Exp Rheumatol. Jan-Feb 2017;35(1):129–136.

58. Jazzar AA, Shirlaw PJ, Carpenter GH, Challacombe SJ, Proctor GB. Salivary S100A8/A9 in Sjogren’s syndrome accompanied by lymphoma. J Oral Pathol Med. Oct 2018;47(9):900–906. doi:10.1111/jop.12763

59. Chen R, Zou J, Liu J, Kang R, Tang D. DAMPs in the immunogenicity of cell death. Mol Cell. Oct 16 2025;85(20):3874–3889. doi:10.1016/j.molcel.2025.09.007

60. Dong H, Zhang Y, Huang Y, Deng H. Pathophysiology of RAGE in inflammatory diseases. Front Immunol. 2022;13:931473. doi:10.3389/fimmu.2022.931473

61. Wang S, Song R, Wang Z, Jing Z, Wang S, Ma J. S100A8/A9 in Inflammation. Front Immunol. 2018;9:1298. doi:10.3389/fimmu.2018.01298

62. Katz J, Stavropoulos F, Bhattacharyya I, Stewart C, Perez FM, Caudle RM. Receptor of advanced glycation end product (RAGE) expression in the minor salivary glands of patients with Sjogren’s syndrome: a preliminary study. Scand J Rheumatol. 2004;33(3):174–178. doi:10.1080/03009740310004775

63. Sreejit G, Flynn MC, Patil M, Krishnamurthy P, Murphy AJ, Nagareddy PR. S100 family proteins in inflammation and beyond. Adv Clin Chem. 2020;98:173–231. doi:10.1016/bs.acc.2020.02.006

64. Leclerc E, Fritz G, Weibel M, Heizmann CW, Galichet A. S100B and S100A6 differentially modulate cell survival by interacting with distinct RAGE (receptor for advanced glycation end products) immunoglobulin domains. J Biol Chem. Oct 26 2007;282(43):31317–31331. doi:10.1074/jbc.M703951200

65. Zhang R, Sun T, Song L, Zuo D, Xiao W. Increased levels of serum galectin-3 in patients with primary Sjogren’s syndrome: associated with interstitial lung disease. Cytokine. Oct 2014;69(2):289–293. doi:10.1016/j.cyto.2014.06.008

66. Uchino Y, Mauris J, Woodward AM, et al. Alteration of galectin-3 in tears of patients with dry eye disease. Am J Ophthalmol. Jun 2015;159(6):1027–1035 e1023. doi:10.1016/j.ajo.2015.02.008

67. Talal N, Grey HM, Zvaifler N, Michalski JP, Daniels TE. Elevated salivary and synovial fluid beta2-microglobulin in Sjogren’s syndrome and rheumatoid arthritis. Science. Mar 28 1975;187(4182):1196–1198. doi:10.1126/science.46621

68. Tecer D, Büyükşireci D, Gunendi Z, Gogus F. The association of serum beta-2-microglobulin with autoantibody production and disease activity in patients with primary Sjögren’s syndrome. Gulhane Medical Journal. 11/23 2020;62:272–277. doi:10.4274/gulhane.galenos.2020.1070

69. Gottenberg JE, Seror R, Miceli-Richard C, et al. Serum levels of beta2-microglobulin and free light chains of immunoglobulins are associated with systemic disease activity in primary Sjogren’s syndrome. Data at enrollment in the prospective ASSESS cohort. PLoS One. 2013;8(5):e59868. doi:10.1371/journal.pone.0059868

70. Menon D, Coll R, O’Neill LA, Board PG. Glutathione transferase omega 1 is required for the lipopolysaccharide-stimulated induction of NADPH oxidase 1 and the production of reactive oxygen species in macrophages. Free Radic Biol Med. Aug 2014;73:318–327. doi:10.1016/j.freeradbiomed.2014.05.020

71. Hughes MM, McGettrick AF, O’Neill LAJ. Glutathione and Glutathione Transferase Omega 1 as Key Posttranslational Regulators in Macrophages. Microbiol Spectr. Jan 2017;5(1)doi:10.1128/microbiolspec.MCHD-0044-2016

72. Menon D, Innes A, Oakley AJ, et al. GSTO1-1 plays a pro-inflammatory role in models of inflammation, colitis and obesity. Sci Rep. Dec 19 2017;7(1):17832. doi:10.1038/s41598-017-17861-6

73. Tan Y, Yin J, Wu Z, Xiong W. Integrative multi-omics analysis reveals cellular and molecular insights into primary Sjogren’s syndrome. Heliyon. Jul 15 2024;10(13):e33433. doi:10.1016/j.heliyon.2024.e33433

74. Harju TH, Peltoniemi MJ, Rytila PH, et al. Glutathione S-transferase omega in the lung and sputum supernatants of COPD patients. Respir Res. Jul 6 2007;8(1):48. doi:10.1186/1465-9921-8-48

75. Piaggi S, Marchi E, Carnicelli V, et al. Airways glutathione S-transferase omega-1 and its A140D polymorphism are associated with severity of inflammation and respiratory dysfunction in cystic fibrosis. J Cyst Fibros. Nov 2021;20(6):1053–1061. doi:10.1016/j.jcf.2021.01.010

76. Pan YC, Chu PY, Lin CC, et al. Glutathione S-transferase omega class 1 (GSTO1)-associated large extracellular vesicles are involved in tumor-associated macrophage-mediated cisplatin resistance in bladder cancer. Mol Oncol. Aug 2024;18(8):1866–1884. doi:10.1002/1878-0261.13659

77. Jiang T, Liu X, Wang S, et al. Paeoniflorin alleviated experimental Sjogren’s syndrome by inhibiting NLRP3 inflammasome activation of submandibular gland cells via activating Nrf2/HO-1 pathway. Free Radic Biol Med. Jun 2025;233:355–364. doi:10.1016/j.freeradbiomed.2025.03.043

78. Khalafalla MG, Woods LT, Camden JM, et al. P2X7 receptor antagonism prevents IL-1beta release from salivary epithelial cells and reduces inflammation in a mouse model of autoimmune exocrinopathy. J Biol Chem. Oct 6 2017;292(40):16626–16637. doi:10.1074/jbc.M117.790741

79. Wei H, Wang JY. Role of Polymeric Immunoglobulin Receptor in IgA and IgM Transcytosis. Int J Mol Sci. Feb 25 2021;22(5)doi:10.3390/ijms22052284

80. Abdelhamid S, Ramirez AV, Aksan E, et al. Dynamic progression of ectopic lymphoid structure formation in lacrimal glands of a Sjogren’s disease murine model. Front Immunol. 2026;17:1797691. doi:10.3389/fimmu.2026.1797691

