## Supplementary_Table_1 for "Tear Proteomics Reveals RAGE and NLRP3 Inflammasome Pathway Activation in Lacrimal Glands of a Sjögren’s Disease Mouse Model"

**Supplemental Table 1: Antibodies and reagents used in Western blotting and Immunofluorescence**

| Purpose | Antibodies/reagents | Vendor | Cat.# | Dilution |
|---|---|---|---|---|
| **Western blotting** | Anti-Galectin-3 | Thermo Fisher Scientific | 14-5301-82 | 1-1000 |
| **Immunofluorescence** | | | | 1-100 |
| **Western blotting** | Anti-S100A8/A9 | Proteintech | 26992-1- AP | 1-1000 |
| **Immunofluorescence** | | | | 1-100 |
| **Western blotting** | Anti-GSTO1-1 | Proteintech | 15124-1-AP | 1-1000 |
| **Immunofluorescence** | | | | 1-100 |
| **Western blotting** | pIgR/SC | House-made | N/A | 1-2000 |
| **Immunofluorescence** | | | | 1-200 |
| **Western blotting** | RAGE | Abcam | ab3611 | 1-1000 |
| **Immunofluorescence** | | | | 1-100 |
| **Western blotting** | NLRP3 | Bioss | bs-10021r | 1-1000 |
| **Western blotting** | Pro-CASP1 | Cell Signalling Technology | 24232T | 1-1000 |
| **Western blotting** | Cle-CASP1 | Cell Signalling Technology | 89332T | 1-1000 |
| **Western blotting** | GSDMD | Cell Signalling Technology | 10137T | 1-1000 |
| **Western blotting** | IL-1β | Abcam | ab283818 | 1-1000 |
| **Western blotting** | IRDye® 680RD Goat anti-Rabbit IgG | LICORbio | 926-68071 | 1-2000 |
| **Western blotting** | IRDye® 680LT Goat anti-Rat IgG | LICORbio | 926-68029 | 1-2000 |
| **Immunofluorescence** | DAPI | Thermo Fisher Scientific | D1306 | 1-1000 |
| **Immunofluorescence** | Phalloidin | Thermo Fisher Scientific | A30107 | 1-200 |
| **Immunofluorescence** | Goat anti-Rabbit-AF488 | Thermo Fisher Scientific | A-11008 | 1-200 |
